# Reproducible social phenotyping of 5xFAD mice in the Agora maze (Sociobox)

**DOI:** 10.64898/2026.02.27.708222

**Authors:** Sheila Sanchez-Garcia, Bettina Platt, Gernot Riedel

## Abstract

Neuropsychiatric (depression, schizophrenia, etc.) and neurological disorders (Alzheimer’s disease, AD, Parkinson’s disease) are characterized by disruptions in cognition including social interaction and recognition. Developing tools for the assessment of social behaviour in mouse models and its relevance is essential to further advance our understanding of social impairments in these diseases. In the Agora maze for rodents, stranger mice confined into cubicles around the perimeter of the open square mirror the agora (marketplace) in ancient cities. Up to 5 social interaction partners are presented and can be freely selected for interaction (exposure). In the discrimination phase one novel mouse (SNew) is presented while 4 familiar partners remain. Interaction time is recorded via video observation.

In Exp 1, we validated the test with different strains of wild-type male mice (C57BL/6J, Balb/c, NMRI) that were able to readily identify SNew and spent significantly more time in zones adjacent to their cubicle; only NMRI mice did not prefer SNew. Exp. 2 explored 5xFAD Alzheimer mice and showed normal exploration and discrimination when aged 6 and 8 months old. Repeat of the experiment in a second cohort confirmed robustness of this phenotype, but also reproducibility of the behavioural paradigm.

The Agora task allows semi-automated evaluation of preference for social novelty in a more complex paradigm by expanding the number of social interaction partners from 2 (three-chamber test) to 5 (or more), while still avoiding physical approaches and aggressive episodes. Thus, Agora provides a more physiological behavioural paradigm which is highly robust and reproducible.

**Highlights:**

- More comprehensive behavioural test bed for social recognition
- Male wild-type mice can identify a stranger mouse amongst 5 social interaction partners
- No deficit in amyloid-based Alzheimer model 5xFAD aged 6-8 months.
- Repeat of experiments returned highly robust and reproducible results.

## 1. Introduction

Human social interaction is crucial to effectively create and maintain relationships in life [1]. Deficits in social behaviour are present in numerous psychiatric and neurological disorders [2] and have a major impact on the quality of life of patients. Alzheimer’s disease (AD) patients for instance display social and cognitive impairments [3] and deficits in face processing [4]. Social isolation is a risk factor of AD [5] and increases the prevalence of neurological diseases [6]. In contrast, slower cognitive decay in socially active individuals during ageing [1,7], thereby also reducing dementia progression. In Parkinson’s disease patients, lower levels of empathy, impaired recognition of facial emotions and frequent dysexecutive behavioural disorders were identified vs. controls [8]. However, mimicking these social behaviours in experimental models have been challenging. Thus, better tools for the assessment of social skills in mouse models are needed to further enrich our understanding of social phenotypes in neurological diseases.

The capacity of rodents to identify a familiar from an unfamiliar conspecific has been well established [9,10]. Social interaction in mice and rats is highly dependent on smell [11]; especially odours emanating from the preputial gland serve as the primary recognition cue in social memory tasks [12]. Tests widely employed are studying social interaction as two-way discriminations, for example in the three-chamber test, also known as Crawley’s sociability and preference for social novelty test [13,14]. In this paradigm the arena consists of three equisized chambers, one on either side containing a small wire-mesh cubicle; chambers are connected to a central compartment by an aperture in the connecting walls. After habituation to the apparatus, the sample trial 1 presents a stranger 1 mouse in the wire-cage on one side, and an empty wire cage on the other. During the test trial 2, a second unfamiliar mouse (stranger 2) is introduced into the empty wire cage, while stranger 1 remains in its familiar position. The idea here is that mice will show a preference towards novelty, i.e. the unfamiliar conspecific stranger 2, and the reduction in interaction with the now familiar stranger 1 may be taken as an index of social recognition [15,16]. Modifications of this apparatus have also been reported [17,18,19,20,21,22]. Using the three-chamber paradigm, scopolamine treated mice showed normal sociability but deficits in social recognition [14]. Similar phenotypes have been reported for genetically modified mouse models of AD [23 and citations therein] but inconsistencies between studies, models and age of subjects remain. Even though this paradigm has proven to be a valuable tool for studying social recognition in wildtype mice as well as for identifying disease-associated social phenotypes, mice are capable of more complex interactions and social behaviours that may be inaccessible with simple 1:1 or 1:2 discriminations. There has now been growing interest in determining more complex social behaviours and novel experimental tools have aided this endeavour ([24] for review). Examples include the development of the ‘Sociobox’ [25], but also more sophisticated analysis tools for social interactions in group housing [26,27,28] or in ethologically relevant home cage-based social recordings of multiple individuals and their social interaction with a single stranger mouse in wire-cages [29, 30]. Especially the latter test conditions, however, are not without issues as home cage cohorts will interact with the stranger in a hierarchically organised manner and the sometimes aggressive socialisation causes highly variable data sets. Moreover, for drug testing it may be difficult to differentiate between animals. We therefore explored a sociobox-like assessment of individual test mice in the presence of multiple stranger mice (more reminiscent of a cocktail party with many unknown guests).

Our term *Agora* is taken from the Greek □γορά, where it was the central place (marketplace) in ancient cities like Athens and Rome, more relevant and translational to human everyday lives. The *Agora* emulates the central square for social activities at the periphery. Consequently, social encounters and active social exchanges are more likely at the perimeter, and the central area is merely an open space. This concept was recently also used by Maurice and coworkers [31] in a more complex ‘Hamlet’ maze.

The experimental design of the Agora maze allows evaluation of preference for social novelty in two ways: 1) the exposure to 5 (or potentially more) strangers arranged around the perimeter of an open square enables detection of preferences for individual stranger mice; 2) the exchange with one (or more) social partner from its cubicle and replacement with a novel stranger mouse (here termed SNew) gives direct comparison of time spent with an unfamiliar conspecific versus time spent with t previously encountered mice. This paradigm therefore provides a more comprehensive assessment of complex social behaviour with avoidance of aggressive episodes. Its complexity is more likely to uncover subtle changes caused by multidimensional diseases, which may be missed by simpler single-cage or 3-chamber arrangements.

We here report on the general setup of a commercially distributed Agora maze (Ugo Basile, Comero, Italy) and on 2 experiments of exploratory research: Exp. 1 compared multiple mouse strains from a commercial breeder and their social behaviour to reproduce the work of Krueger-Burg et al [25]. Exp. 2 quantified the social interaction of the 5xFAD model of Alzheimer’s disease (AD) at various ages against wild-type controls. Despite previous evidence of social interaction deficits in this AD line, we (and others) failed to identify impairments in social cognition [32, 33]. By testing of several cohorts, we also monitored the robustness and reproducibility of the paradigm.

## 2. Materials and methods

### 2.1. Animals

Subjects for Exp. 1 were male wild-type animals of the same sex and strain (C57BL/6J, Balb/c, PLB_WT_ (wild-type) or NMRI mice purchased from Charles River (n=10 each) aged 12-15 weeks at the beginning of testing. PLB_WT_ were derived from C57BL/6J mice (Charles River Laboratories CRL UK) and crossed with C3H mice more than 100 generations ago to generate the PLB1_Triple_ AD mouse [34]. They were maintained by backcrossing with C57BL/6J mice and are expected not to differ from standard C57BL/6J strain from the distributor. As stranger mice we used male C57BL/6J (CRL UK) aged 10-15 weeks at the beginning of testing, housed in a different part of our animal facility.

For Exp. 2, male 5xFAD transgenic mice (C57BL/6J background) and wild-type littermate controls were used. Five x FAD mice express 5 mutations previously associated with familial AD including APP K670N/M671L (Swedish), I716V (Florida), V717I (London), and presenilin 1 M146L + L286V mutations (B6.Cg Tg (APPSwFlLon, PSEN1*M146L*L286V; 6799Vas/Mmjax, JAX MMRRC Stock# 034848) and were derived from crossing heterozygous transgenics with wild-type C57BL/6J mice in our local animal facility and maintained until aged 5 months in cohorts of mixed genotypes. All other conditions were as in Exp. 1. Ear biopsies were analysed for genotype identification for all offspring by Transnetyx Inc. (Cordova, USA) and hAPP/PS1 positive male mice were selected together with non-transgenic littermates.

The two replications of Exp. 2 were performed using 15 heterozygous 5xFAD (familial Alzheimer’s disease: replication 1 n=7; replication 2 n=8) and 15 wild-type littermates (replication 1 n=7; replication 2 n=8) male mice aged 20 weeks at the beginning of testing.

Replication 2 was performed several months after replication 1 and is thus treated as independent. Since we were unable to find a deficit in social behaviour in 5-month-old animals, animals of replication 2 were aged further until 8 months old and were again tested in the Agora. Experimentally naïve C57BL/6J (CRL, UK) served as strangers. Male mice were selected because social interactions are less rewarding for female mice [35] and they previously failed to show recognition in the sociobox [25]. However, in the 3-chamber apparatus, wild-type female mice could socially discriminate [36].

Mice were group-housed in standard Macrolon II size cages (corncob bedding with cardboard tubes and paper wool as enrichment) within different rooms for the test and the stranger animals. They were classed as Specific Pathogen Free (Exp. 1) at arrival from Charles River Laboratories by truck and acclimatised for at least 1 week to the holding facility. Maintained in open cage housing, animals had free access to food (Special Diet Services, Witham, UK) and water at an ambient room temperature of 21 ± 1◦C and 40–55% relative humidity. A day-night cycle of 12 hours (light on: 7am) with simulated dusk and dawn (30 mins) was implemented; animals were checked daily and cleaned once per week.

Mice were tested in accordance with UK Home Office regulations (project licence number PP2213334). All experimental procedures were subject to the University of Aberdeen’s Ethics Review Board (AWERB) and conducted in accordance with the European Directive on the Protection of Animals used for Scientific Purposes (2010/63/EU) and the Animal (Scientific Procedures) Act 1986 and its amendments from 2012 and in accordance to the ARRIVE 2.0 guidelines [37].

### 2.2. Agora apparatus

The *Agora* apparatus was provided by Ugo Basile SRL (Comero, Italy) as a modification of the method previously published by Krueger-Burg D et al [25] termed sociobox. Made of grey Perspex, it consisted of a decagonal open arena in the centre (diameter = 36 cm, height = 21 cm) and five external cubicles (11.5×13 cm) situated around its perimeter equidistant between each other (Fig. 1 for details) into which stranger mice were placed. Cubicles contained a front facing transparent plastic slider with holes (∼2 mm in diameter) up to 9.5 cm of height. These holes permitted the exchange of odours and sound between animals during social interaction. All walls were removable to facilitate the cleaning of cubicles between tests.

**Figure 1.**
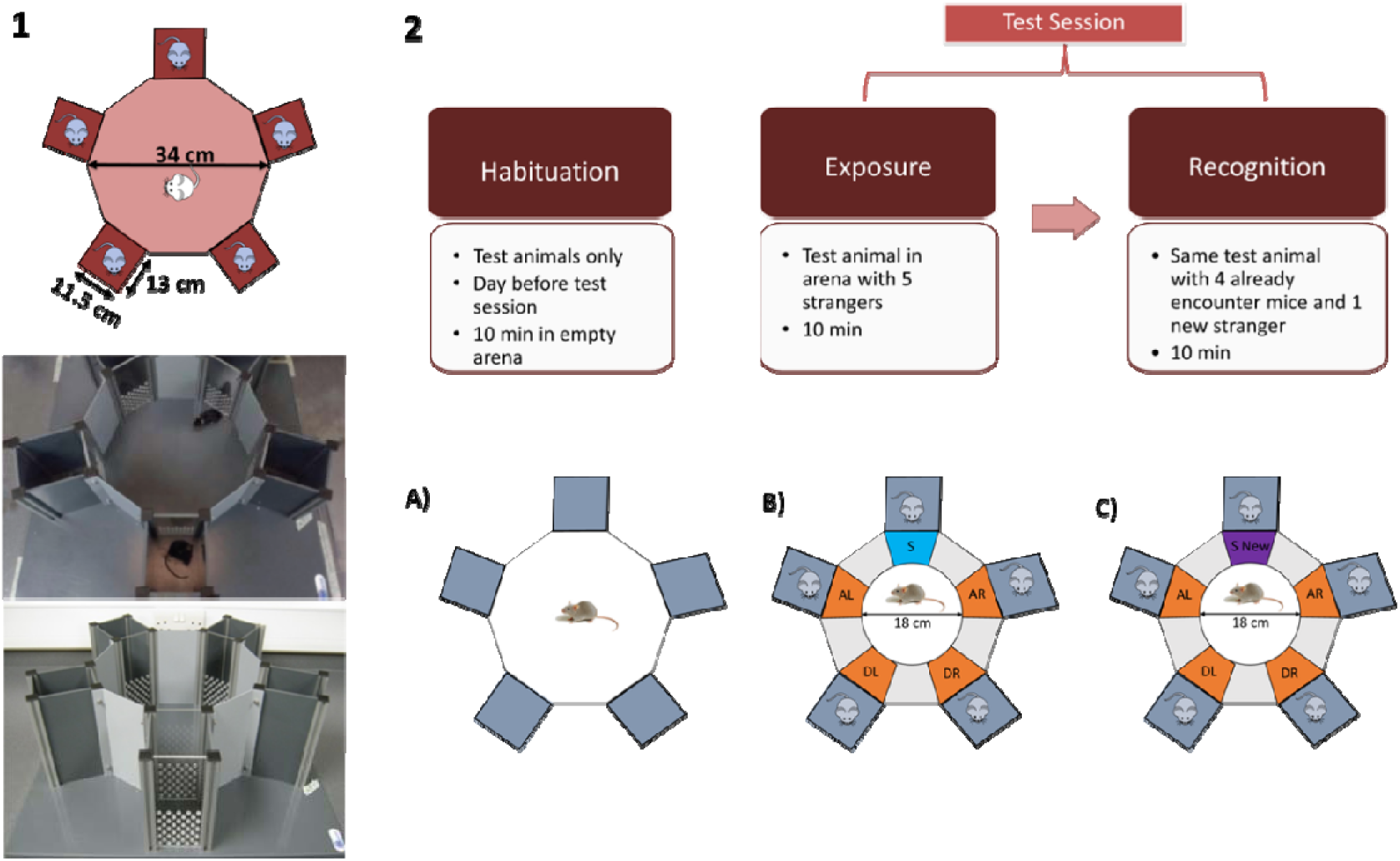
Agora apparatus and experimental design. (**1**) Diagram of the apparatus incl. dimensions and photographs of the arena with examples of mice interacting. (**2**) Experimental outline of 3 stages including details for habituation (**A**) exposure (sampling) (**B**) and recognition (discrimination) (**C**) phases of the test. Experimental details of each phase are also denoted. The diagrams show the 5 equally sized zones adjacent to the strangers that have been used for the analysis of the data (S, stranger; SNew, stranger new; AL, adjacent left; AR, adjacent right; DL, distal left; DR, distal right).

### 2.3. Video tracking

Behaviour was recorded with an overhead CCTV camera (Sanyo VCB-3385P) fixed ∼100 cm above the arena. The camera was connected to a PC controlled video card and recorded using the tracking software ANY-Maze (Ugo Basile SRL, Varese, Italy; Version 5.26) at 10Hz sampling rate. Tracking was attained using back-ground subtraction with darker or brighter fur colour of test subjects. Homogenous illumination of the arena was achieved by indirect lighting (∼30 Lux).

### 2.4. Experimental setup

All experiments were conducted during the light phase (9 am to 5 pm). Animals (test and stranger mice) were acclimatized to the experimental room for 30 min at the beginning of each session. The running order was randomised, but experimenters were not blinded and no power calculations were performed *a priori*. Group sizes were taken from experience with other social interaction work conducted previously using the 3-chamber apparatus [14,34,38,39,40].

#### 2.4.1. Habituation session

On day 1, each test animal was habituated to the central arena; fronts of the cubicles were shut but cubicles themselves remained empty. Test mice were released into the centre of the decagonal arena and allowed to freely explore the maze for 10 minutes, before being returned to their home cages. All parts of the arena were cleaned after each animal with a wet towel and dried using paper towels.

#### 2.4.2. Test session

On day 2, five stranger mice were introduced into the cubicles at the outer perimeter of the Agora maze. The test session consisted of two phases: *exposure* and *social recognition* (Fig. 1). For the exposure phase the test animal was placed into the central area and left to freely explore the arena and socialize with stranger mice for 10 mins. At the end of this phase the test animal was returned to its transport cage, the Agora maze was cleaned (see above) and for the recognition phase one stranger mouse was replaced by a new stranger (SNew); all other animals remained in the same position as during the sampling phase. After an inter-trial interval of 2-3 minutes, the test animal was placed back into the arena and freely explored the Agora for another 10 mins. Several sets of stranger mice were used during the experiments to avoid social disinterest and discomfort. Furthermore, to avoid any spatial bias, the position of SNew was rotated by 1 position clockwise for each test animal.

The primary data analysis was performed using ANY-Maze. Five zones of equal size adjacent to the cubicles were defined as social interaction zones (11 x 16 cm) and the time spent (and number/duration of visits) in these zones by the test animals was automatically calculated as a proxy for social interaction with any stranger mouse (Fig. 1). This was contrasted with time in the contact zone of SNew during recognition. A recognition index (ΔT) was also calculated as interaction time with SNew minus the average of interaction time spent with the other 4 stranger mice. Other parameters analysed included distance travelled and speed for the Agora as well as time in outer (16 cm) and inner zones (18 cm).

### 2.5. Statistical analysis

Statistical analyses were performed using the Prism software V10 (GraphPad, CA, US). Outlier detection followed the method of Grubbs with a significance level of α= 0.01. Data were confirmed as Gaussian and analysed using repeated measures one-way Analysis of Variance (ANOVA) or repeated measures two-way ANOVA (with genotype as between-subject and time as within-subject factor) followed by post-hoc t-tests with Bonferroni correction; two-group comparisons were performed using two-tailed unpaired Student t-tests. Data are displayed as group Mean +/- SEM with scatter of individual data points; alpha was set to p<0.05. Only significant outcomes are denoted in the Results for simplicity.

## 3. Results

### 3.1. Exp. 1: Social behaviour varies between wild-type mouse strains

We here compared 4 different wild-type mouse strains as test mice: C57BL/6J, PLB WT, Balb/c, and NMRI. Habituation to the Agora presented an open field-like exploration test and we recorded locomotor activity [41] in different zones (outer and inner) as index of anxiety-related behaviour [42].

During *habituation* the total distance travelled in the Agora on day 1 differed between strains (F(3,34) = 3.327; p = 0.031), but this did not apply to both the exposure and recognition phases of the test (F’s < 1.7; p’s > 0.18). Activity differences were primarily due to lower values for Balb/c mice relative to all other lines (p < 0.05; Fig. 2A). We also recorded the distribution in the periphery and confirmed reliable differences for stages (F’s > 2.6; p’s < 0.02) habituation (Fig. 2B) and recognition (Fig. 2H) but not for exposure (Fig. 2E). A heightened anxiety is indicated for the periphery (thigmotaxis zone) preferring NMRI mice. Exploration of the centre also yielded several differences especially in the PLB_WT_ cohort, which ventured into this zone more often when stranger mice were presented (F(3,34) = 9.7; p < 0.0001 for exposure, F(3,34) = 5.2; p = 0.0045 for recognition; Figs. 2F & I).

**Figure 2.**
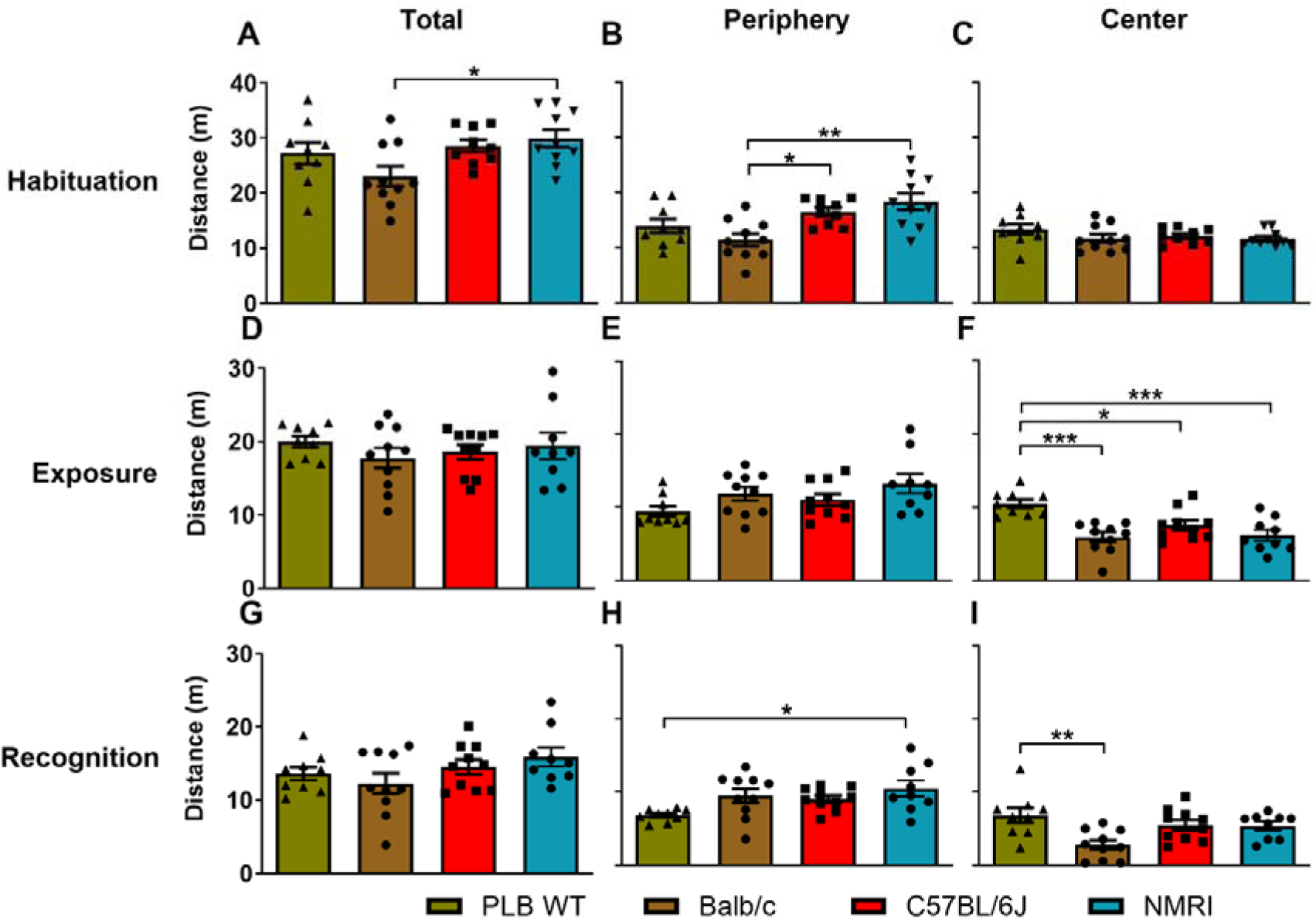
Differences in activity between strains during Habituation and Test sessions (exposure and recognition). Total distance travelled by the mice in the Agora (left column), the wall zone (periphery – middle column) and in the center (right column). No strangers were present during habituation. Small differences were observed for total activity (**A,D,G**). It appeared that NMRI mice travelled more in the periphery than other strains (**B,H**), while PLB_WT_ mice (as their sister strain C57BL/6J) ventured more into the centre (**F,I**). Statistical analysis was conducted using one-way ANOVA and Bonferroni post-hoc test. *p<0.05, **p<0.01, ***p<0.001. Data are presented as Mean +/- SEM plus individual data as scatter.

Not unexpected was a global drop of activity in all strains during the *exposure* phase once stranger animals were placed in their cubicles (distal left, DL; adjacent left, AL; adjacent right, AR; distal right, DR). The activity shifted to the interaction zones as part of the periphery due to sociability with their peers (compare Figs. 2A, D & G for global activity and Fig. 2B, E, H for activity in the periphery close to the stranger mice). At the same time, there was less exploration of the centre (Fig. 2C, F &I). This difference between periphery and centre was particularly prominent for the Balb/c cohort but not seen in C57BL/6J and PLB_WT_ mice.

We further segmented the exploration of the cubicles by selective measurement of the time spent in the *interaction zones* in close proximity of each stranger during exposure (Fig. 3A-D) but did not find any bias for a specific individual in any of the lines (all F’s<1). During recognition, when one mouse was replaced by SNew, there was a significant main effect of location in all lines (F’s > 2.8; p’s < 0.04, Fig. 3I-K) apart from the NMRI cohort which failed to show a spatial bias (F(4,24) = 1.6; p = 0.2; Fig. 3L). The strongest recognition was expressed by the Balb/c mice (F(3,33) = 5.9; p = 0.0025; Fig. 3M) as is also confirmed from the heat maps of the radar plots (Fig. 3E-H). We also calculated ΔT as the difference of time spent with SNew relative to the mean of time spent with all 4 familiar mice (Fig. 3N) as an index for the strength of preference. The positive values in PLB_WT_, Balb/c and C57BL/6J mice indicate a strong bias for SNew; yet this was not observed in NMRI mice. This suggests a lack of recognition memory for strangers in the NMRI cohort, an observation coupled with increased propensity to escape the apparatus during the experiment in NMRI male mice. None of the other strains showed such behaviours.

**Figure 3.**
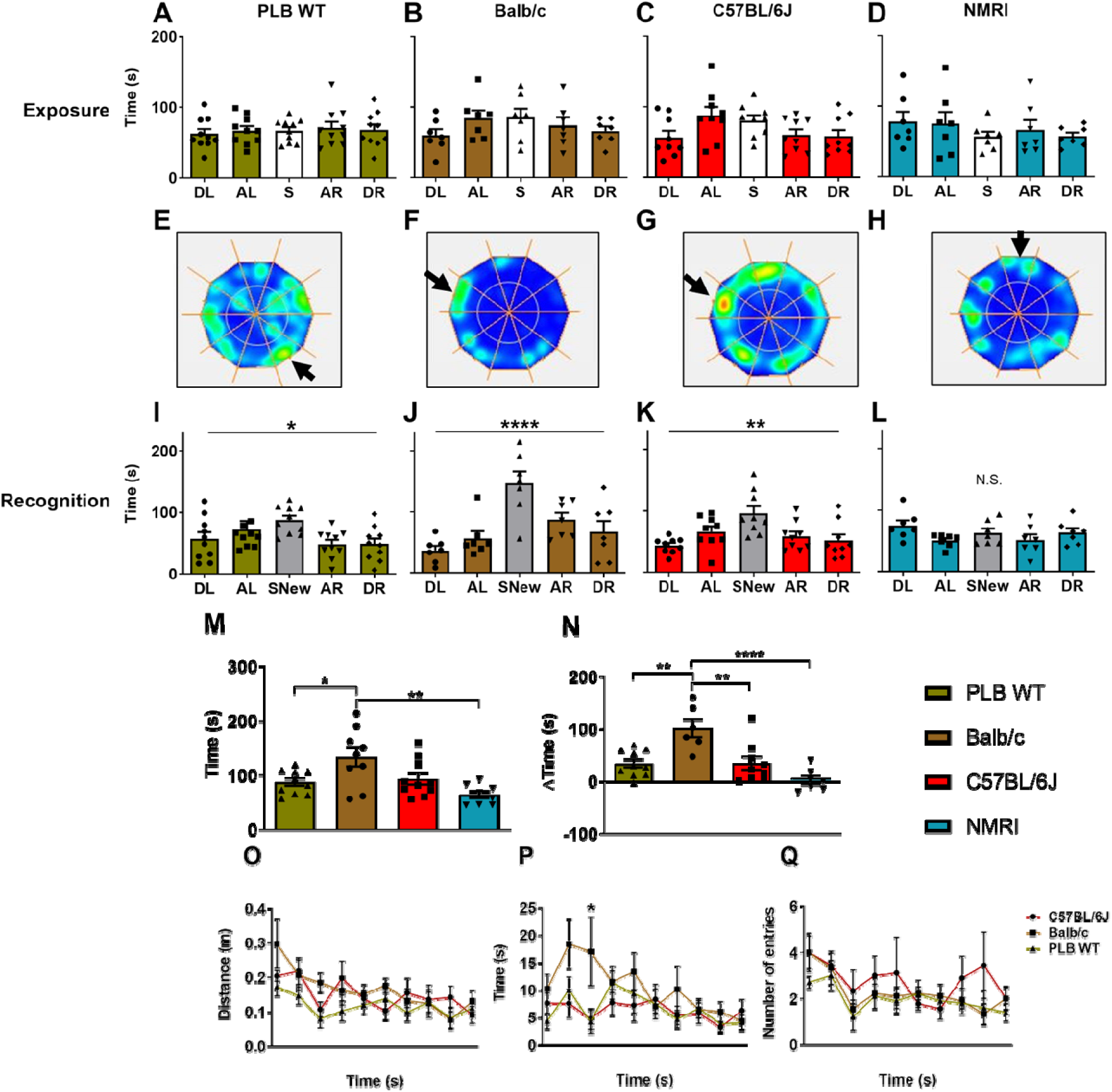
Performance of wild-type lines during exposure (sampling) and recognition (discrimination) phases in the Agora maze. A-D: Time spent in the interaction zones with stranger mice for the different strains. No spatial bias was observed for any strain. The open bar indicates the stranger mouse to be replaced for recognition. For abbreviations, see below. **E-H:** Representative radar plot heat maps of the activity of individual test mice during the recognition test: PLB_WT_ (E), Balb/c (F), C57BL/6J (G), NMRI (H). Arrows indicate location of SNew (randomised within groups), and warmer colours reflect higher amounts of time spent in this zone. All animals show preference for the interaction zone of SNew. **I-L:** Quantitative data for time spent in the interaction zones during recognition testing. All strains apart from NMRI mice (**L**) showed a significant difference for time in zones with a bias for the zone of SNew (grey bars). **M**: Strain comparison of time spent with SNew; Balb/c mice outperformed all other strains. **N**: Social recognition index (DT) for all wild-type strains. Discrimination was best in Balb/c, but there was no recognition in NMRI mice. **O-Q:** Time course for visits to interaction zone SNew (NMRI excluded) during recognition: Distance travelled by the test mice inside zone SNew. **(O);** Average bout time of each visit (**P**); Number of visits (**Q**). Animals showed heightened levels of activity in early stages of the session, activity wanes towards the end. Balb/c mice presented with longer visiting bouts after release (1-4 minutes). Statistical analyses were conducted using repeated measures one or two-way ANOVAs with post-hoc t-tests. Data are expressed as Mean +/- SEM including individual data as scatter for bar charts. *p<0.05, **p<0.01, ***p<0,001. Abbreviations: (S, stranger; SNew, stranger new; AL, adjacent left; AR, adjacent right; DL, distal left; DR, distal right).

For a more in-depth analysis of the performance during the recognition phase, we analysed the time course of the distance travelled inside the zone SNew (Fig. 3O), the average duration per visit (Fig. 3P) and the number of visits (Fig. 3Q). For all parameters, we obtained a significant effect of time (F’s > 2.5; p’s < 0.006) indicative of habituation to SNew with interest waning over the 10-minute period. No group differences were observed for distance moved, but there was a significant main effect of strain for both number of visits and time spent in the zone per visit (F’s > 3.6; p’s ≤ 0.04). Only Balb/c mice expressed prolonged visits (Bonferroni: Balb/c vs. C57BL/6J p = 0.0255, Balb/c vs. PLB_WT_ p = 0.0312).

### 3.2. Exp. 2: No deficit in social novelty or social recognition in 5xFAD, but highly reproducible outcomes

After validation of the Agora maze in male wildtype strains, we moved on to confirm sensitivity of the test, and to determine the existence of a social phenotype using the 5xFAD amyloid model for Alzheimer’s disease (AD). Animals are known to rapidly develop severe amyloidosis and high levels of intraneuronal Aβ42 already around 1.5 months of age.

Amyloid deposition emerges in around two months old mice, first in the subiculum and layer 5 of the cortex and increasing in density and spreading with age. Plaques are developed in hippocampus and cortex by six months of age. Synapse degeneration is also observed at approximately four months coincident with neuronal loss and deficits in spatial learning at approximately four to five months [43,44,45,46].

We initially tested 5 months old 5xFAD and their wild-type littermates (N=7 each; replication 1). A second cohort (N=8; replication 2) was run a few months later through the same protocol in order to measure robustness and replicability of the paradigm and the transgenic mice. Globally, we were able to reproduce the data from cohort 1 exactly with cohort 2 and did not find any statistical differences for the factor replication. As already established for the different wild-type strains, ambulatory activity was highest during habituation and decreased while animals were in exploration and recognition phases (cohort 1: Fig. 4A-C; cohort 2: Fig. 4D-F). There was no genotype difference in either cohort. Similarly, the time spent in the respective interaction zones with the 5 stranger mice during exploration was not different between genotypes or replication (cohort 1: Fig. 4G, H; cohort 2: Fig. 4I, J) and there was no spatial bias for any single stranger mouse (F’s < 1.7; p’s ≥ 0.19). As shown in the radar plots and the quantitative data (cohort 1: Fig. 4K, L, O, P; cohort 2: Fig. 4M, N, Q, R), a strong bias for SNew was revealed for all groups in the recognition phase (F’s > 3.5; p’s ≤ 0.02) with a near perfect replication of the cohort 1 data in cohort 2. Similarly, no differences were found between genotypes and replications for the recognition index (ΔT) for cohorts 1 (Fig. 4S) and cohort 2 (Fig. 4T). These data confirm robustness of the paradigm and its reproducibility in wild-type and 5xFAD mice.

**Figure 4.**
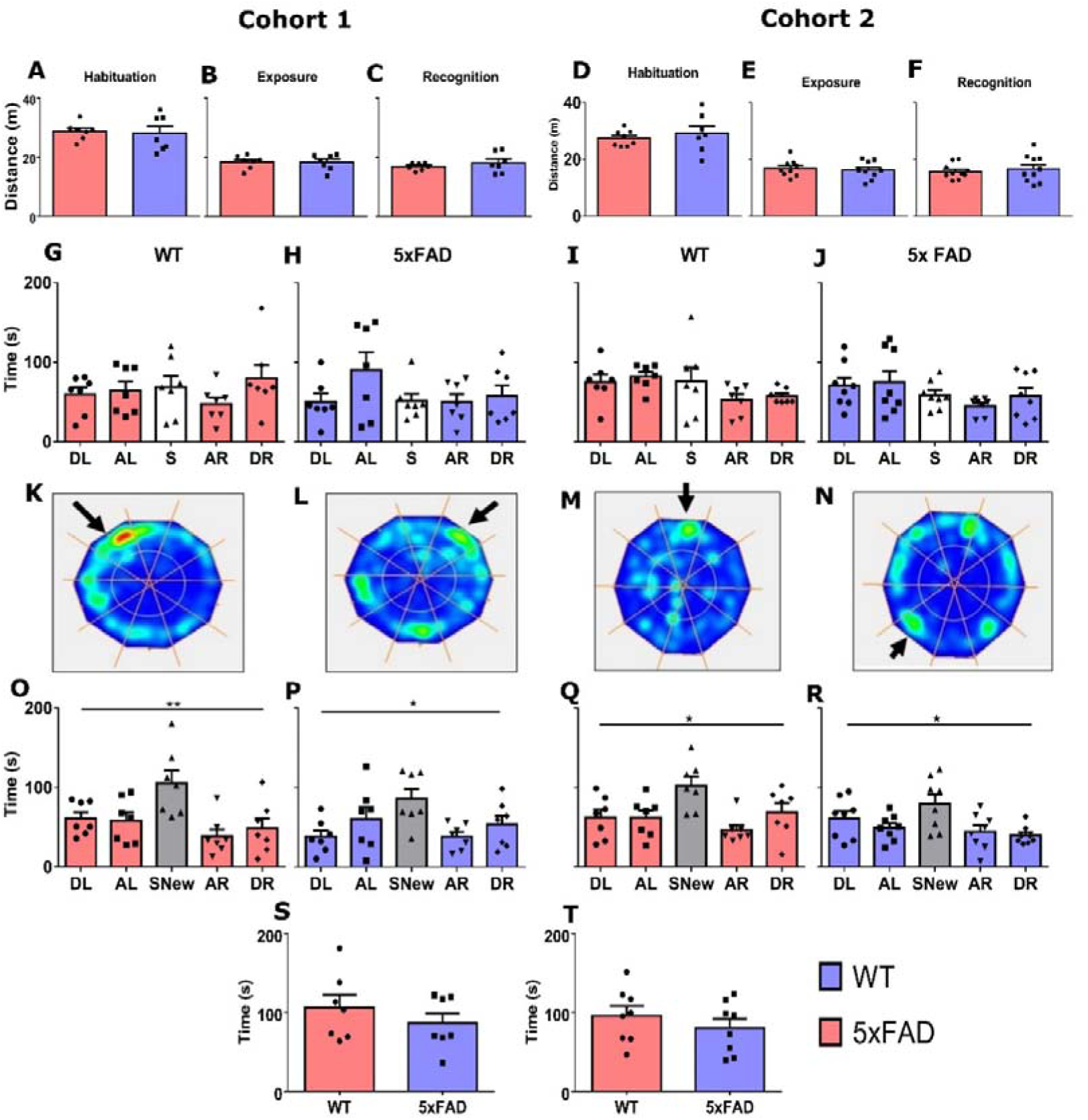
Social interaction in two cohorts of wild-type and 5xFAD male mice aged 5 months. Test for reproducibility. A-F: Total ambulatory activity did not differ between genotypes for any of the test phases. Data were reliably reproduced between replications in cohorts 1 and 2. **G-J:** Interaction times with 5 stranger mice during the exposure phase. There was no spatial or social bias for any one of the stranger mice and no effect of genotype. Cohort 2 **(I,J)** reproduced the data of cohort 1 **(G,H).** Open bars indicate stranger to be replaced with SNew. **K-N**: Radar heatmaps of representative performances during recognition testing. Arrows indicate location of SNew (randomised within groups) and warmer colours reflect higher amounts of time spent in this area. All animals show preference for the interaction zone of SNew. **O-R**: Quantitative data for time spent in the interaction zones during recognition testing. Both genotypes/cohorts showed a strong bias for the zone of SNew (grey bars). Data were highly reproducible. **S,T**: Genotypic comparison of time spent by the test animals in the interaction zone of SNew for cohort 1 (**S**) and cohort 2 (**T**). Both replications returned the same results, i.e. no phenotype in 5xFAD mice. Statistical analysis employed repeated measures one-way ANOVAs followed by student t-tests if appropriate. Data are expressed as Mean + SEM and individual data are shown as scatter. *p<0.05, **p<0.01.

This lack of phenotype at 5 months of age replicated our previous work using 5xFAD and PLB1_Triple_ mice in barrier based social interaction protocols [32,34]. Yet others have suggested social deficits in older amyloid models (see [47] for review) or specifically in 5xFAD mice [49]. We thus aged cohort 2 until 8 months old and repeated the test with a new set of age-matched stranger mice. Total distance travelled during habituation was drastically reduced at 8 months in both genotypes relative to the same cohort at younger age (Fig. 5A). A two-way ANOVA returned a main effect of age (F(1,17) = 251.6, p < 0.0001) and an interaction between age and genotype (F(1,17) = 13.95, p = 0.0016). Post-hoc t-test with Bonferroni correction confirmed the reduction in ambulatory activity in both genotypes but also returned heightened activity levels in 5xFAD mice relative to wild-type (Fig. 5A). During the test session, 8-month-old wild-type and 5xFAD mice did not different in social recognition (Fig. 5B recognition index). Their social contacts also did not differ during the exploration phase (Fig. 5C, D) and both genotypes presented with discrimination of SNew (Fig. 5E – wild-type: F(4,28) = 3.483, p = 0.0198; Fig. 5F - 5xFAD: F(4,32) = 2.974, p = 0.0339).

**Figure 5:**
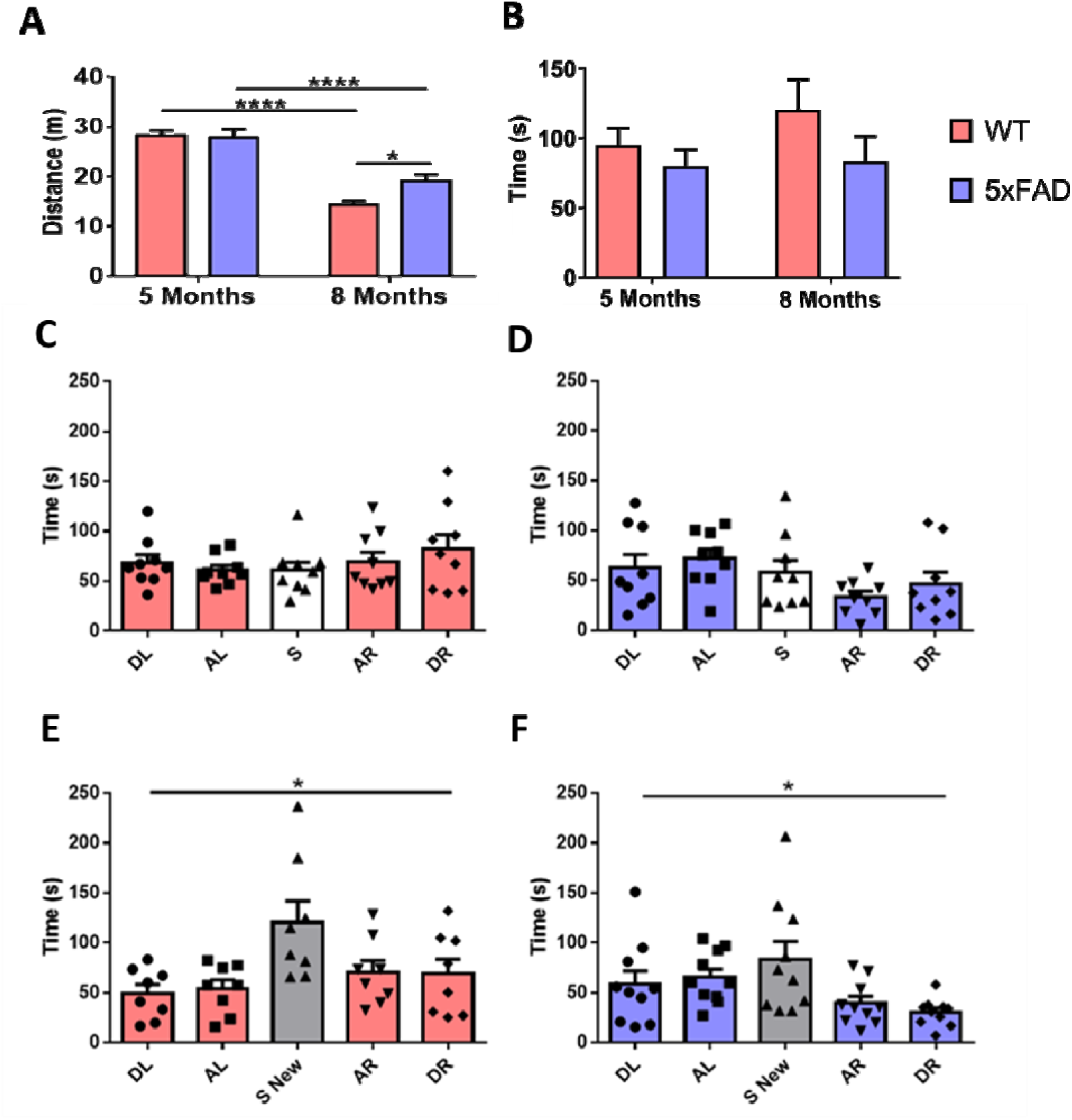
Agora test session data for 8-month-old 5xFAD and wild-type littermates. **A:** Comparison of ambulatory activity during habituation for 5 and 8-months-old cohorts. A reduction of activity with age was observed in both genotypes; yet 5xFAD mice were hyperactive at 8 months relative to age-matched wild-types. **B**: Comparison of time spent in interaction zone with SNew during recognition for 5 and 8-month-old cohorts. There were no differences between genotypes for either replication. **C, D**: Interaction times with 5 stranger mice during the exposure phase at 8 months of age. There was no spatial or social bias for any one of the stranger mice and no effect of genotype. Open bars indicate stranger to be replaced with SNew. **E, F**: Quantitative data for time spent in the interaction zones during recognition testing. Both genotypes showed a preference for SNew (grey bars; asterisks indicate significant difference between locations (1-way ANOVA)). Statistical analysis employed one-way ANOVAs followed by student t-tests if appropriate. Data are expressed as Mean + SEM and individual data are shown as scatter. *p<0.05, ****p<0.0001.

## 4. Discussion

### 4.1. Validation of the Agora with different wildtype strains of mice

Social interactions and social recognition are important skills to effectively create and maintain relationships in life and lack of normal social exchanges is a key characteristic of many neurodevelopmental and neurodegenerative diseases. Developing tools for the assessment of social skills in rodent models is essential to further advance our understanding for its mechanistic underpinnings. Here, we evaluated the validity of a complex social recognition paradigm, which we termed the Agora maze, and in which stranger mice are confined to cubicles around the perimeter of a central arena, that is occupied by a test subject. In Exp. 1, we assessed behavioural differences in various wild-type strains (C57BL/6J, Balb/c, PLB_WT_, and NMRI) in a mixed confirmatory/exploratory study. We confirmed heightened activity to novelty in C57BL/6J and NMRI compared with Balb/c mice [49,50]. Contrary to previous reports indicating high anxiety levels of Balb/c mice in paradigms such as the open-field, light-dark box and elevated plus maze [49,51,52], Balb/c mice displayed no more thigmotaxis in this paradigm than all other strains. Yet, differential ambulatory activity has been reported between various wild-type mouse strains for other arenas [53,54], which are most likely due to the arena format and the recording setup (see [55] for differences due to recording equipment).

The preference for novelty in rodents has been well established [1 and citations therein] and indicates that the reduction in interaction with the previously encountered mouse in favour of an unfamiliar stranger SNew may be taken as a proxy for social recognition. Our data corroborate findings of Krueger-Burg and co-workers [25] for C57BL/6J and Balb/c that these strains, to which we add NMRI, are differentially able to discriminate SNew among five possible interaction partners. Despite low levels of ambulatory activity, Balb/c mice showed the best recognition for SNew while NMRI mice did not seem to identify the novel stranger (see Fig. 3N). Interestingly, there were no gross performance differences between the two C57 sublines (C57BL/6J and PLB_WT_) indicating that many years of independent breeding did not affect this behavioural trait. Observations in the 3-chamber apparatus with exposure to two interaction partners revealed that male C57BL/6J, DBA/2J and FVB/NJ strains did not differ in social recognition [13,36]. The difference here is that the Agora presents a more complex and naturalistic approach with high translational validity compared to simpler single cage and 3-chamber tests. A more substantial study using the Agora with all strains present awaits completion.

Another widely reported observation is a reduction in overall activity of the test animals during exposure and recognition compared to habituation, also shown here [14,36,56]. The introduction of stimulus mice clearly presents a focus for the test mouse and while the wire-cages in the 3-chamber or single chamber arrangement allow free access from all angles, the interaction in the Agora is one directional. Nevertheless, the animals in the Agora also focus on the interaction partners (see Fig. 3E-H) and do not reduce their activity in the outer periphery during the different test stages. All mouse strains tested are behaving similarly. Differential outcomes were reported for the 3-chamber box [56,57] showing that Balb/c mice expressed a much greater reduction in activity during sociability and social recognition. Although not explicitly tested here, we suggest that performance in the Agora is driven by intra rather than extra maze cues. Our experimental design counterbalanced the position of SNew within each group, and we did not find preferences for any specific spatial orientation. However, since context has been suggested to play a major role in social interaction [58] this requires more rigorous testing. Similarly, it is unclear whether the heighted rcognition index in male Balb/c mice (this study; [59]) is correlated with their superior level of agression [60,61].

### 4.2. Lack of impairment in social recognition in the Agora in 5xFAD mice is highly reproducible

After validating the experimental set up with wildtype strains and given our interest in neurodegenerative models, we tested an Alzheimer’s disease model which carries 5 mutations previously associated with familial AD (5xFAD). This experiment had two purposes: i) we sought to determine whether there are deficits in social behaviour in male 5xFAD mice aged 6 months, and ii) to test the reproducibility and robustness of their performance in the Agora. Consequently, two cohorts were assessed with several months in-between experiments. The overarching outcome was that 5xFAD mice were not deficient in sociability or social recognition at this age, and that the test was highly robust and reproducible in both wild-type C57BL/6J and transgenic 5xFAD (Fig.4).

Deficits in social withdrawal widely reported in Alzheimer’s disease patients [3,4,62] have been confirmed in various rodent experimental disease models ([14,34,65] but see [66]). However, controversial results have been obtained for 5xFAD mice: some reports mentioned no deficits at ages of ≤ 6 months [23,32,67,68,69] while others reported deficits in 6 months or older cohorts (6 m – [70]; > 10 m – [23,71]). It therefore appears that the social phenotype develops with age and is more robustly observed in male mice, despite a heightened amyloid load in female 5xFAD mice [32]. Whether a correlation can be established between Aβ40/42 ratios and social cognition was not investigated here but would be an interesting research subject for future experiments. Since we had previously failed to determine social memory deficits in the OF-NO-SI (open field, novel object, social interaction) apparatus, in which only a single mouse is presented in a wire mesh cage and the habituation between two trials is recorded as a proxy of recognition 32,40], we reasoned that a complex arrangement with more interaction partners could reveal social deficits. Nevertheless, even in animals up to 8 months of age, we failed to observe a social interaction and recognition deficit in 5xFAD mice. By contrast, Nguyen and co-workers [66] reported different results from two replications of the same experiment using 3xTg-AD mice. This prompted a similar approach here to determine the robustness of our outcome. Data in 6-month-old mice turned out to be highly robust and reproducible due to exact replication of experimental conditions within the same laboratory. We and others have previously shown that maintaining identical experimental conditions are an important prerequisite for data reproduction in animal behaviour even between laboratories [55,72,73,74,75]. We here add to the list of behavioural tests used in the above studies the social interaction test Agora, which appears as a highly translationally relevant approach to determine social cognition.

Since we did not find deficits in 5xFAD mice, it is unlikely that animals up to 8 months of age developed severe impairments to odour cues that provide primary sensory input in social memory tasks [12] although such deficits have been previously acknowledged [76,77,78]. Similarly, apathy-like neuronal processes are not impaired given normal or hyperactivity in 8-month-old 5xFAD animals, despite evidence that such phenotypes appear in such middle-aged cohorts [33].

In conclusion, the Agora paradigm produced robust social recognition in wild-type mice. Male mice can readily identify a stranger mouse (SNew) and spent significantly more time in a zone adjacent to its cubicle compared to the several previously encountered conspecifics. This paradigm provides a more realistic behavioural scenario than other commonly used social recognition tests and offers the possibility of determination of more complex parameters, such as comparison between recent and remote memory of multiple social partners.

### Institutional review board statement

All animal experiments were performed in accordance with the European Communities Council Directive (63/2010/EU) with local ethical approval of the University of Aberdeen (AWERB) under the UK Animals (Scientific Procedures) Act (1986; amended in 2012), a project licence (PP2213334) and complied with the ARRIVE guidelines 2.0. No human samples were used in this study.

## CRediT authorship contribution statement

**Sheila Sanchez-Garcia:** Study planning, Methodology, Investigation, Formal data analysis, Visualization, Data curation, Writing – original draft. **Bettina Platt:** Supervision, Resources, Writing - review & editing. **Gernot Riedel:** Conceptualisation, Supervision, Resources, Project administration, Funding acquisition, Writing – review & editing,

## Data Availability

Data will be made available on request.

